# Oolonghomobisflavans from *Camellia sinensis* disaggregate tau fibrils across Alzheimer’s disease models

**DOI:** 10.1101/2024.02.26.582120

**Authors:** Chatrawee Duangjan, Xinmin Chang, Paul M. Seidler, Sean P. Curran

## Abstract

Alzheimer’s disease (AD) is a common debilitating neurodegenerative disease with limited treatment options. Amyloid-β (Aβ) and tau fibrils are well-established hallmarks of AD, which can induce oxidative stress, neuronal cell death, and are linked to disease pathology. Here, we describe the effects of Oolonghomobisflavan A (OFA) and Oolonghomobisflavan B (OFB) on tau fibril disaggregation and prionogenic seeding. Transcriptomic analysis of OF-treated animals reveals the induction of a proteostasis-enhancing and health-promoting signature. OFA treatment reduced the burden of Tau protein aggregation in a *C. elegans* model expressing pathogenic human tau (“hTau-expressing”) and promoted Tau disaggregation and inhibited seeding in assays using *ex vivo* brain-derived paired helical filament tau protein fibrils from Alzheimer’s disease brain donors. Correspondingly, treatment with OF improved multiple fitness and aging-related health parameters in the hTau-expressing *C. elegans* model, including reproductive output, muscle function, and importantly, reversed the shortened lifespan stemming from pathogenic Tau expression. Collectively, this study provides new evidence supporting the neuroprotective effects of OFs and reveal a new therapeutic strategy for targeting AD and other neurodegenerative diseases characterized by tauopathy.

## INTRODUCTION

Age-associated neurodegenerative diseases are a major public health challenge, due to recent increases of lifespan and the absence of effective pharmacological treatments [1]. Alzheimer’s disease (AD) is the most common neurodegenerative diseases characterized by cognition and memory impairments [1]. The typical markers of AD histopathology are the accumulation of amyloid β-protein (Aβ) and neurofibrillary tangles (NFTs) in brain tissue which drive neuronal dysfunction and cell death [1, 2].

Tau is a microtubule-associated protein that interacts with tubulin to promote and maintain microtubule stability [2]. Under physiological conditions, tau participates in the regulation of microtubule assembly, which impacts axonal transport and the structural organization of the synapse [1, 2]. In tauopathies, pathological transformation of tau begins with hyperphosphorylation, conformational changes of protein structure, loss of microtubule-binding affinity, oligomerization, misfolding and ultimately the formation of insoluble filaments that accumulate as neurofibrillary tangles NFTs [1, 2]. Hyperphosphorylation of tau further promotes the formation of proteotoxic intracellular amyloid aggregates that impact neurodegenerative diseases [2]. The loss of microtubule stability due to abnormal of tau phosphorylation has been reported as a major cause of tauopathies [2].

*Caenorhabditis elegans* (*C. elegans*) has been extensively used as a model of neurodegenerative diseases [3]. Transgenic *C. elegans* strain KAE112, has been created with codon-optimized human 0N4R V337M tau expressed in the body wall muscle to better understand the impact of pathogenic tau expression on cellular function [4]; hereafter referred to as hTau-expressing model. In this model, the hyperphosphorylation of the human tau variant drives proteotoxicity, resulting in animals that display premature defects in age-associated health metrics, including reproductive fitness, developmental rate, muscle paralysis, and even lifespan [4]. Increasing evidence suggests the development of interventions that target tau could be potent treatments for AD and other tauopathies [5]. Therefore, the *C. elegans* hTau-expressing model is optimal for designing therapeutic strategies and developing drug screening methods for anti-tauopathy treatments.

Several studies have reported oxidative stress and neuronal cell damage as key drivers of AD pathology [6]. Natural products from herbs or plant extracts with potent antioxidants that inhibit tau aggregation could provide an alternative approach to treat or prevent neurodegenerative diseases. Tea polyphenols act as natural bioactive compounds that could complement traditional therapeutic agents for neurodegenerative diseases characterized with proteostasis defects [7, 8], including Aβ [9-11], tau [12], α-synuclein [10], inflammation [13] and oxidative stress [9-11].

Oolong tea (*Camellia sinensis*) has been studied for beneficial effects on neurodegenerative diseases [13, 14], but the molecular mechanisms underlying the neuroprotective effects of bioactive compounds in oolong tea require further investigation. In previous work, we identified the oxidative stress resistance properties and neuroprotective effects against Aβ of oolong tea extracts and its bioactive molecules oolonghomobisflavans (OFs) [9-11]. In this study we further identified the specific action of oolonghomobisflavan A (OFA) and oolonghomobisflavan B (OFB) on tau fibrils aggregation and tau protein-induced toxicity by using *C. elegans* and human Alzheimer’s brain extract models, which suggests OFs might be a valuable therapeutic strategy for the development of new nutraceutical preparations that target neurodegenerative diseases.

## RESULTS

### OFA and OFB treatment induce a proteostasis-enhancing and health-promoting transcriptional signature

Based on our documented health and longevity effects of OFA and OFB (OFs) on WT animals and amyloid models of proteinopathy [9], we wanted to understand the molecular basis of this response. To that end, we first compared the transcriptional profile of WT with and without OFs treatment by RNAseq. 511 mRNA transcripts are significantly different (including 194 up-regulated and 317 down-regulated) with OFA treatment when compared with untreated controls (**Figure 1A** and Table S1), 660 mRNA transcripts (including 261 up- and 399 down-regulated transcripts) are significantly different in OFB treatment when compared with untreated controls (**Figure 1B** and Table S1), and 72 mRNA transcripts (including 69 up- and 3 down-regulated transcripts) are significantly different in OFA treatment when compared with OFB treatment (**Figure 1C** and Table S1).

**Figure 1.**
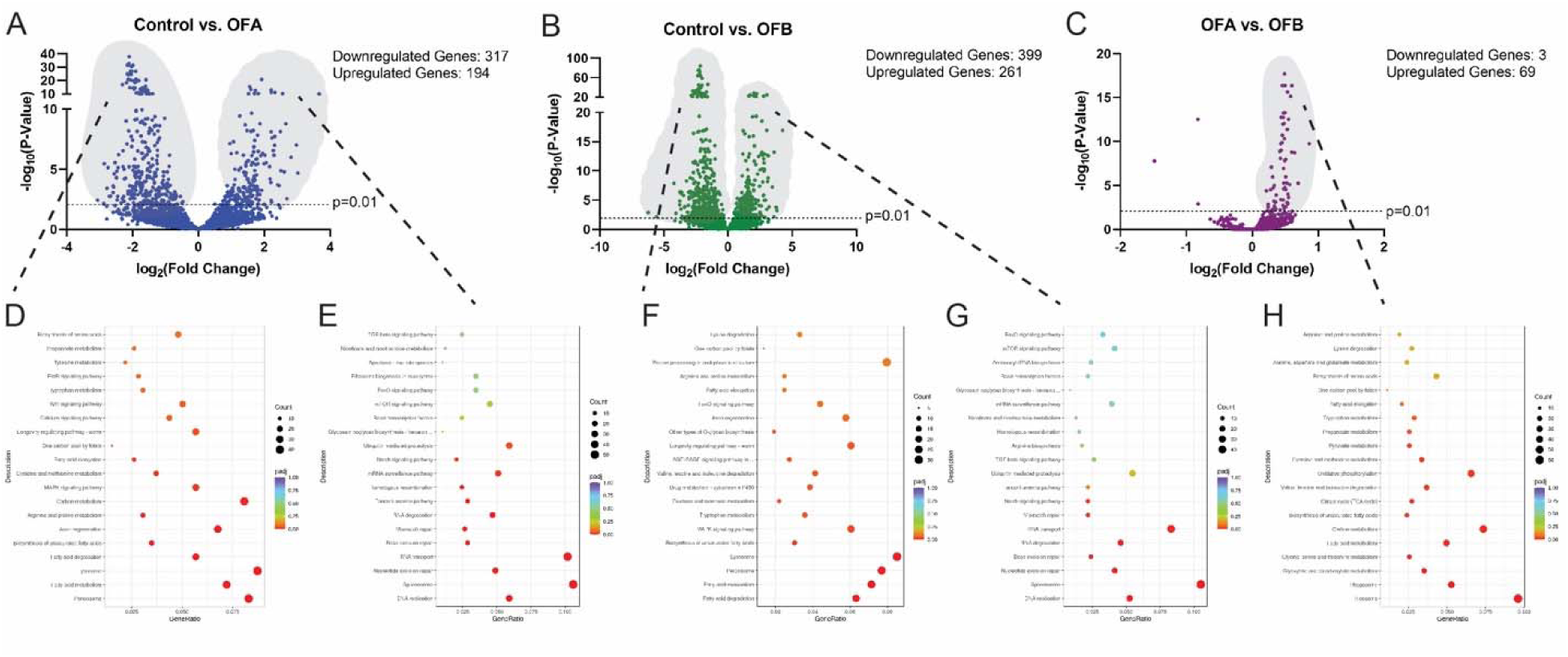
OF treatment induces a healthspan promoting transcriptional signature. Volcano plots of differentially expressed genes between mock-treated controls and OFA (**A**) and OFB (**B**) and genes differentially expressed between OFA and OFB treatment (**C**). As compared to mock treatment (controls), gene ontology (GO) and KEGG enrichment analysis of genes that decrease (**D,F**) and increase (**E,G**) expression in OFA and OFB treatment, respectively. (**H**) GO and KEGG enrichment analysis of genes that increase expression between OFA and OFB treatment. The mean expression level for each gene is indicated by log2FoldChange. All genes considered to be significant has adjusted p-value < 0.05.

We performed a gene enrichment analysis (GEA) for each treatment group and identified classes of transcripts significantly regulated by OFs (**Figure 1D-H**, Figure S1A-G, and Table S1). Several oxidative stress-related terms were identified, including response to stress (GO:0006950), cellular response to stress (GO:0033554), cellular response to DNA damage stimulus (GO:0006974), oxidation-reduction process (GO: 0055114), and oxidoreductase activity (GO:0016491). KEGG analysis revealed enrichment for longevity-regulating pathways (KO:04212 and KO:04213); including), FoxO (KO:04068), mTOR (KO:04150), and MAPK (KO:04010) signaling pathways. Importantly, among the genes differentially altered between OFA and OFB treatment was the expression of several ubiquitin-mediated proteolysis (KO:04120 and GO:0016579), Rho and RAS signaling (GO:0017048 and GO:0017016), and axon regeneration (KO:04361) genes were regulated by OFA treatment; albeit near significance for OFB. Taken together, these results demonstrate that OF treatment influences the expression of genes that promote oxidative stress responses and cellular proteostasis and signaling that help maintain homeostasis and that OFA and OFB treatment have remarkably similar impact on the transcriptional landscape.

### OFs treatment disaggregates human tau fibrils from AD brain extracts

In light of the measured enhancement of cellular proteostasis responses in animals treated with OFA and OFB, we next investigated the neuroprotective effects of OFs on animals expressing pathogenic human tau variants (0N4R;V337M), hereafter called “hTau-expressing”, which can lead to shortened lifespan and diminished health [4].

In early stages of Alzheimer’s disease, tau becomes hyperphosphorylated and mislocalized, which can contribute to its aggregation and toxicity [15, 16]; and this hyperphosphorylation is mimicked in the *C. elegans* hTau-expressing model [4]. To measure the impact of OFs treatment on tau protein dynamics we examined the phosphorylation status (**Figure 2A-C**) and aggregation propensity (Figure S2A) of the hTau protein in the hTau-expressing animals treated with OFA and OFB as compared to mock-treated controls. The abundance of phosphorylated Tau (P-tau) on residue S202 was reduced ∼81-82% and S416 was reduced 78-88% and the abundance of aggregated Tau, as measured by the slower migrating population in the polyacrylamide gel, was also significantly reduced. Taken together these results reveal that OFA and OFB influence Tau proteostasis.

**Figure 2.**
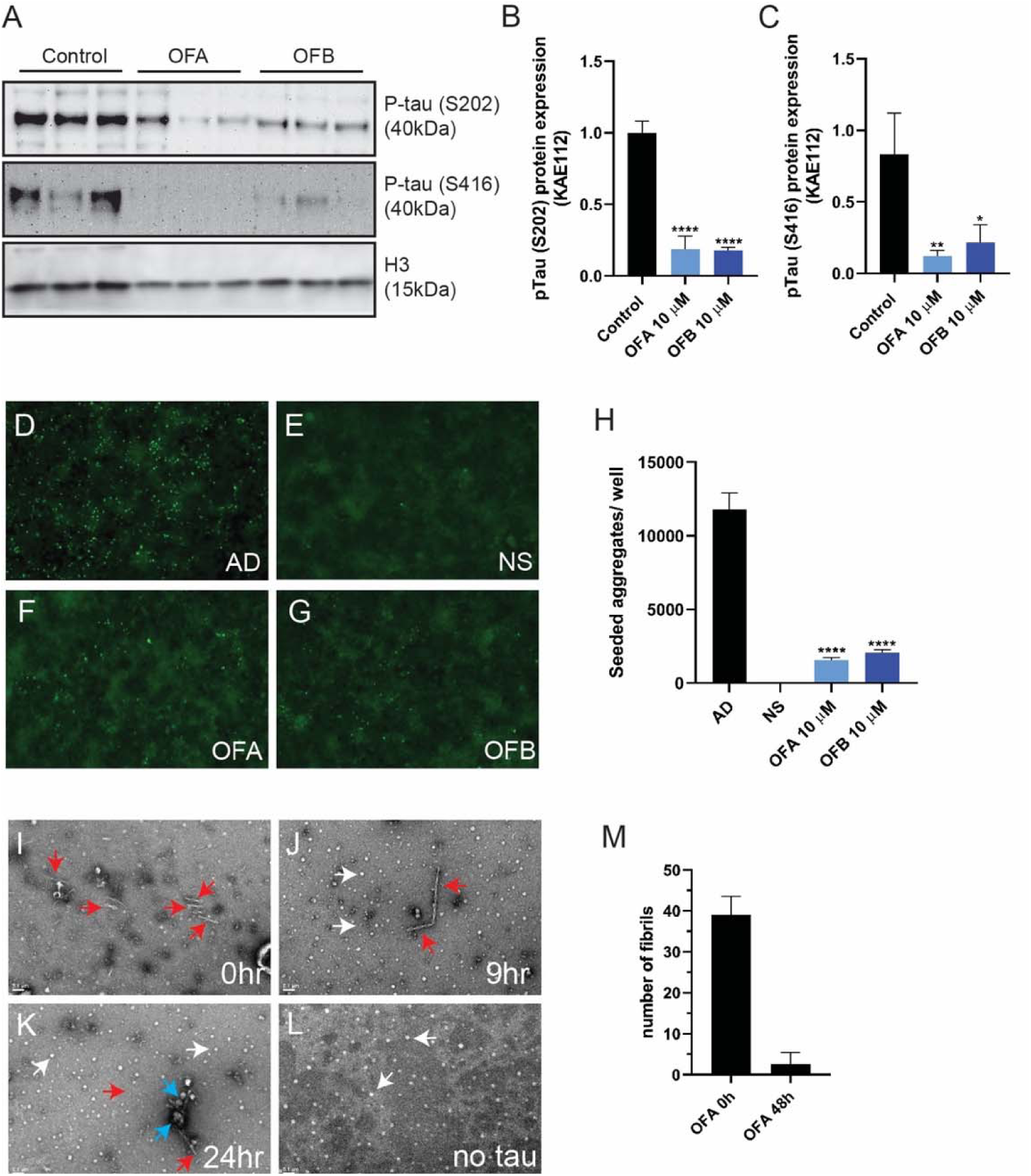
OF treatment reduces tauopathy across model systems. (**A-C**) Western blot analysis of tau phosphorylation (S202, S416) in hTau-expressing worms (KAE112) worms after treatment with OFA and OFB relative to mock-treatment (conrol). (**D-H**) Seeding inhibition measured by transfecting inhibitor-treated AD brain homogenate in fluorescent tau K18 biosensor cells; (**D**) AD brain homogenate without added inhibitor, (**E**) NS, No Seed, (**F**) OFA, (**G**) and OFB. (**H**) Seeded aggregates were determined by quantifying the number of fluorescent puncta as a function of indicated inhibitor and disaggregation. Error bars represent standard deviations of triplicate measures. (**I-M**) OFA-mediated AD tau fibril disaggregation, measured by qEM of AD tau fibrils. (**I-L**) Representative images shown of AD tau fibrils (red arrows), OFA condensates (white arrows), and OFA-associated fibrils (blue arrows). (**M**) Fibrils quantified after 24 hr incubation with OFA. Fibrils were counted from randomly acquired micrographs obtained by automated imaging using EPU software from N=66 images. Fibril counts were obtained by splitting the image sets three ways. Error bars represent standard deviations. *p<0.05, **p<0.01, ***p<0.001 and ****p<0.0001, compared to the mock-treated controls by one-way ANOVA following Bonferroni’s method (post hoc).

Tau aggregation results in the formation of fibrillary structures that propagate and drive neurodegeneration by prion-like seeding, that can spread from one cell to others [2]. We next examined whether OFs treatment could inhibit tau fibril formation in a biosensor cell assay that can measure fibril disassembly capability and proteopathic seeding activity [12, 17]. We observed the dose-dependent inhibition of tau seeded aggregation by crude AD brain homogenate in both OFA and OFB treatment groups, as compared to mock treated controls (**Figure 2D-H**).

We next investigated the capacity of OFs to promote the disaggregation of brain purified AD-tau fibrils by using quantitative electron microscopy (qEM) [12, 17]. AD brain-derived tau fibrils were incubated with OFA for 0, 3, 9, and 24h at 37°C (**Figure 2I-L** and Figure S2B-D). As compared to the abundance of tau fibrils (red arrows) present at the start of the assay (0 h) by 3 hours of incubation, OFA condensates (white arrows) become more pronounced with instances of condensates coalescing with AD tau paired helical filament (PHF) fibrils (blue arrows) (Figure S2B). By 9 hrs, tau fibrils lose their fibril-like morphology suggesting disaggregation by OFA treatment (**Figure 2J** and Figure S2C). At 24 hrs, the presence of tau fibrils is significantly reduced, and fibrils that remain are largely engulfed by OFA condensates (blue arrows) (**Figure 2K** and Figure S2D). Images quantified after 24 hrs of treatment with OFA shows a ∼95% reduction in AD tau PHFs (**Figure 2M**). These data suggest that oolonghomobisflavins are a potent bioactive molecules capable of disaggregation of human brained derived AD-tau fibrils.

### OFs treatment reverses physiological detriments of tauopathy in *C. elegans*

The ability OFs to reduce tau proteinopathy both *in vivo* and *in vitro*, suggested that OFs would also be able to alleviate the health-related detriments stemming from expression of hTau. Because we previously noted a general improvement in health with age in animals treated OFs, we next characterized the effects of OFs in detail in both WT and hTau-expressing animals. Previous studies have documented the negative effects of tau proteotoxicity on multiple fitness parameters, developmental growth, and timing and brood size [4]. OFA and OFB treatment were both capable of significantly reversing the impaired reproductive output of hTau-expressing worms (**Figure 3A-B**), specifically towards the end of the reproductive span at days 3-5 of adulthood (Figure S3A). In contrast, OFA and OFB treatment had no effect on the slowed development and growth observed in hTau-expressing animals (**Figure 3C-D**).

**Figure 3.**
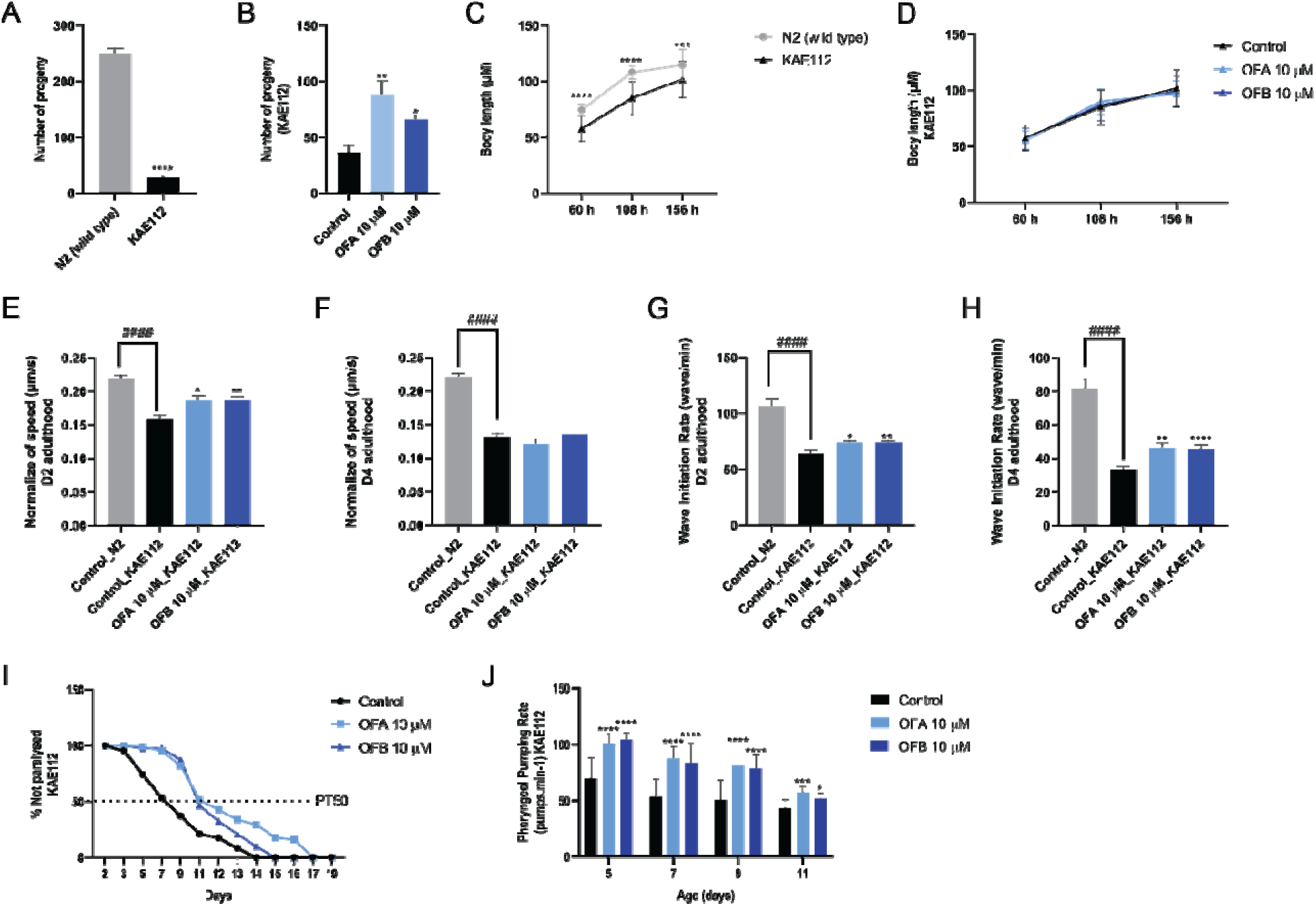
Effects of OFs on health metrics of in hTau-expressing animals. Comparison of total viable progeny (**A,B**), and body size (**C,D**) between wild type (N2) and hTau-expressing worms (KAE112) worms treated with OFA and OFB, as compared to mock treatment (control). Effect of OFA and OFB on crawling speed (**E,F**), thrashing (**G,H**), movement paralysis (**I**) and pharyngeal pumping rate (**J**) in wild type (N2) and hTau-expressing worms (KAE112) worms, as compared to mock treatment (control).*p<0.05, **p<0.01, ***p<0.001 and ****p<0.0001, compared to the mock-treated controls by one-way ANOVA following Bonferroni’s method (post hoc).

We next measured the impact of OFA and OFB treatment on the decline in muscle function resulting from hTau-expression [4]. We confirmed the impaired crawling and thrashing speed in hTau-expressing animals and noted a significant improvement of both movement parameters with treatment of either OFA or OFB at Day 2 and Day 4 of adulthood. By Day 4, thrashing speed showed significant improvement with treatment, while crawling speed did not exhibit significant changes (**Figure 3E-H** and Figure S4AB). Moreover, hTau-expressing worms display premature body-wall and pharyngeal muscle paralysis, as measured by the time where 50% of the worms are paralyzed, but this early decline of age-associated cellular impairment is significantly delayed by ∼4 days in animals treated with OFs as compared to the mock-treated control group (**Figure 3I**).

Pharyngeal function is a facile biomarker of aging in *C. elegans* [18], which displays accelerated decline in hTau-expressing animals (Figure S4C). We previously demonstrated that OF treatment could protect pharyngeal function with age in WT animals [9] and similarly, pharyngeal pumping rate was significantly improved with OF treatment in hTau-expressing animals on days 5, 7, 9, and 11 of adulthood as compared to mock-treated controls (**Figure 3J** and Figure S4C).

### Oolonghomobisflavans reverse shortened lifespan of *C. elegans* model of tauopathy

We confirmed the shortened lifespan previously documented in animals expressing pathogenic human tau (**Figure 4A)** and found that wild-type (N2) and hTau-expressing worms (KAE112) treated with OFA and OFB at the L4 larva stage displayed a significant extension of lifespan in both genotypes as compared to mock-treated controls (**Figure 4B-C**). In general, co-treatment with both OFA and OFB (OFAB) did not provide and synergetic effects (Figure S5AB) which suggests OFA and OFB extend lifespan by similar mechanisms as predicted by the remarkably similar transcriptional profiles documented (**Figure 1**).

**Figure 4.**
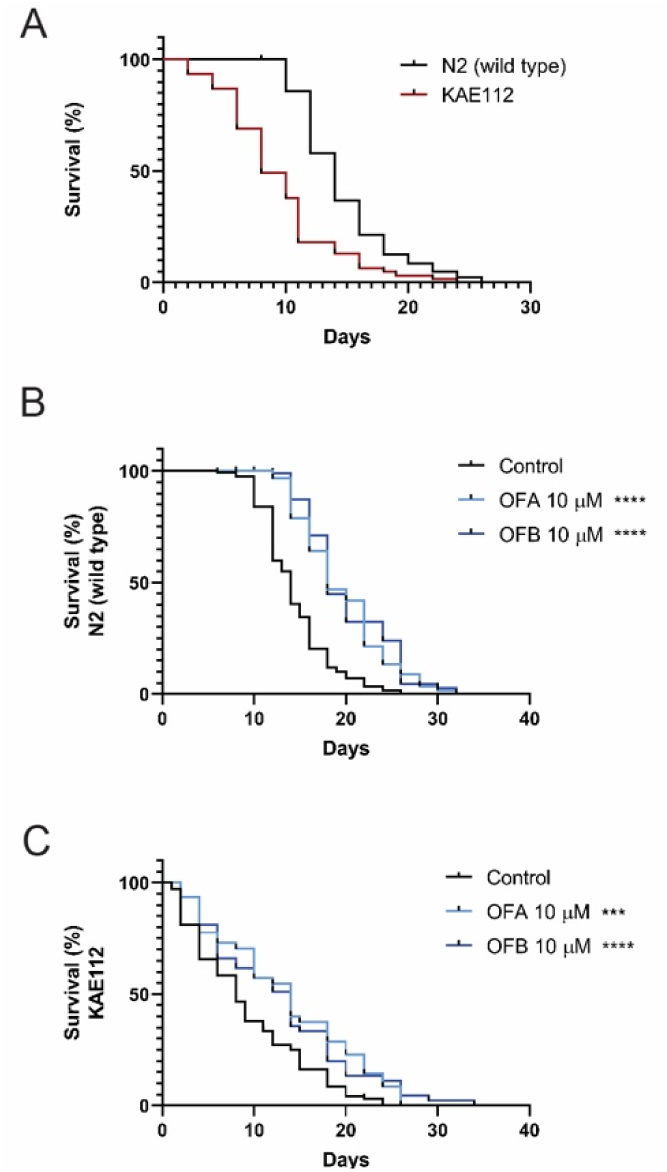
Effects of OFs on lifespan of KAE112. **(A)** Comparison of lifespan between wild type (N2) and hTau-expressing worms (KAE112). Survival curves of wild-type (N2) **(B)** and hTau-expressing worms (KAE112) (**C**) worms treated with OFA and OFB. *p<0.05, **p<0.01, ***p<0.001 and ****p<0.0001, compared to the mock-treated controls by one-way ANOVA following log-rank test.

Collectively, these data reveal that treatment with OFs can drive a lifespan promoting enhancement of organismal health, but more importantly can significantly delay the age-related dysfunction in cells with hTau-related proteotoxicity.

## DISCUSSION

In this study, we demonstrate the ability of OFs to ameliorate tau proteotoxicity across tauopathy models. Our results reveal mechanistic insights underlying the neuroprotective and longevity-promoting effects of OFs that can guide the development of treatments targeting tauopathy and other neurodegenerative conditions.

Oolonghomobisflavans are structurally distinct dimers of the tea polyphenol, epigallocatechin gallate (EGCG) that have emerged as potent antioxidant agents that can contribute to overall neuronal health [7, 12]. Multiple lines of evidence suggest that EGCG has the capacity to impact tauopathy [12, 19-21], but the mechanistic details of these effects require additional investigation. In this study, we illuminated the effects of OFs on tauopathy at the organismal-level and reveal the capacity for OFA to disaggregate tau fibrils extracted from post-mortem brains of AD patients. Electron micrographs suggest that OFA forms condensates, which functionally interact with and disaggregate tau fibrils. The aggregation of chemically similar polyphenols has been characterized at high resolution [22] and is driven by phenol-phenol interactions. Condensates of OFA are suited to maintain productive phenol-mediated binding to tau fibrils given the excess number of aromatic hydroxyls (N=16) for OFA and OFB.

The discovery of molecules that can treat AD has been challenging, but compounds that are able to interfere with the transformation process, which is catalyzed by prionogenic seeding by tau fibrils could be a promising prophylactic or therapeutic strategy for neurodegenerative diseases. Our results demonstrate that OFs can reverse tau aggregation in *C. elegans* and lysates from human AD brains, but also promote health improvements in the *C. elegans* tauopathy model. *C. elegans* worms which expressed human tau exhibit a shortened lifespan, due to tau-induced proteotoxicity, proteostasis loss, apoptosis and eventually cause death [3, 4, 23]. Similar to the lifespan-promoting effects of OFs on WT animals, OFs treatment can also increase the lifespan of the short-lived tau-expressing *C. elegans* model of tauopathy. Our data define the molecular underpinnings of this longevity response, particularly the enhancement of oxidative stress resistance, maintenance of DNA damage, and perhaps most importantly, the enhancement of proteostasis and dissolution of tau aggregates that follows treatment with OFs.

Although strikingly similar, we noted some minor differences between the transcriptional profiles of OFA and OFB treated animals. *In vitro*, there is little difference in the capacity of OFA or OFB to inhibit seeding by tau fibrils, but these differences *in vivo* might be linked to the variation in organismal-level responses to OFs. It has been demonstrated that DAF-16/FoxO transcription factors are necessary for protective effects against the Aβ and tau-induced proteotoxicity in *C. elegans* [24, 25]. Intriguingly, several genes in DAF-16/FoxO signaling pathway were up-regulated in OFs treatment, including: PI3K/AKT(*age-1, akt-2, sgk-1, aap-1*), AMPK (*aak-2, aakb-1,aakg-1, cyb-1, cyb-2.1, cyb-2.2, cyb-3*), p38MAPK/MAPK (*mpk-1, pmk-1, pmk-3, plk-1, par-4, jnk-1*), PI3K/AKT (*age-1, akt-2, sgk-1*), and FoxO (*daf-3, daf-2, daf-9, daf-12, daf-18, sir-2.1, age-1, nhr-80, mtl-1*). These data suggest that DAF-16/FoxO may be targeted by OFs as a mechanism, whereby OFs may alleviate tauopathies.

Oxidative stress is closely related to age-related neurodegenerative diseases [1]. In previous studies, we demonstrated the oxidative stress resistance properties and neuroprotective effects against Aβ of oolong tea and OFs in cultured neuronal cells and *C. elegans* models [9-11]. In this work, we found several oxidative stress-related genes, including catalase (*ctl-1, ctl-2*), glutathione peroxidase (*gst-4, gst-6*), glutathione dehydrogenase (*gsto-1*), and ShkT-containing peroxidase (*skpo-1, skpo-3*) were up-regulated by OFs treatment. In addition, collagen genes including *rol-6, sqt-1,* and *nas-33* were up-regulated in OFs treatments, which have been previously reported to influence neurodevelopmental toxicity and oxidative damage [26].

DNA damage is a well-established driver of the natural aging process that can accelerate the pathogenesis of neurodegenerative diseases [27]. High levels of oxidative stress has been demonstrated to drive DNA damage in the hippocampal tissue in normal aging and AD patients, suggesting a correlation of DNA damage and pathological changes in AD [27]. We found genes related to cellular response to DNA damage-induced stress including *mre-11, pmk-3, pxn-2, ercc-1, polh-1, pif-1, msh-4, hpr-9, nth-1, cku-80, crn-1, lig-1,* and *tipn-1* were upregulated by OFs treatment. Notably, Mre11 complex protein plays an important role in cellular stress responses to DNA damage and has been found to be substantially reduce in the cortical neurons of AD patients [28]. In this study, we found *mre-11* was upregulated by OFs treatment suggesting DNA maintenance as a potential target of OFs to alleviate tauopathies. Further studies to examine oxidative stress resistance effects of OFs on tau-mediated oxidative stress and DNA damage responses is needed.

Importantly, we document the ability of OFs treatment to increase expression of several ubiquitin mediated proteolysis genes including ubiquitin-specific proteases (*usp-3,usp-4, usp-39, usp-46, usp-48*), and deubiquitinase ( *duo-1, duo-3, cyld-1*), Rho/Ras GTPase signaling (*pix-1, unc-73, rhgf-1*), and axon regeneration genes (*rhgf-1*, *pmk-3* and *pxn-2*) [29, 30]. Beyond the clear role that the UPS plays in proteostasis, the GTPase signaling pathway plays a critical role in neuronal functions. The Rho GTPase family regulates actin and microtubule dynamics, while the RAS GTPase family acts as membrane-associated signal transducers that control neuronal survival and regeneration, synaptic connectivity, growth, and differentiation [31]. In addition, Rad23, a nucleotide excision repair, plays a role in protein degradation and has been reported in proteolysis of TNFs through polyubiquitination in the proteasome [32]. The observed up-regulation in OFs treatment, suggest Rad23 as a potential target of OFs, supporting the inhibition effect of tau-inducing proteotoxicity in vivo and disaggregation of brain-derived tau fibrils in vitro.

One clear class of molecules up-regulated by OFs that can contribute to the enhanced proteostasis are the heat shock protein (HSP) class of chaperones, which can mitigate protein misfolding, aggregation, and accumulation [33]. The disruption of proteostasis is associated with the progression of amyloid plaques and tau tangles in neurodegenerative diseases [33]. The chaperone-associated activities of the Hsp family of proteins, like Hsp16.2, that regulates protein folding and maintains homeostasis by reducing the aggregation and toxicity of Aβ and tau in across model systems [22, 23]; Hsp27 which reduces neuronal tau accumulation and restored hippocampal plasticity [34], and Hsp22 that can inhibit heparin-induced aggregation and tau protein levels in cultured cells [35]. Collectively, the enhancement of the cellular proteostasis machinery is a likely contributor of the ability of OFs to maintain proteostasis even in the presence of pathogenic tau expression.

Taken together, our study leverages two complementary tauopathy models which reveal OFs as a potent new class of molecules that can be leveraged to oppose the age-related accumulation of protein aggregation that can lead to disease. These results suggest OFs could be well suited as interventions to counteract systemic proteinopathies. Further studies in *C. elegans* and other models of neurodegeneration are needed to evaluate organismal effects of OFs on other disease states in the CNS.

---

**Figure S1.**
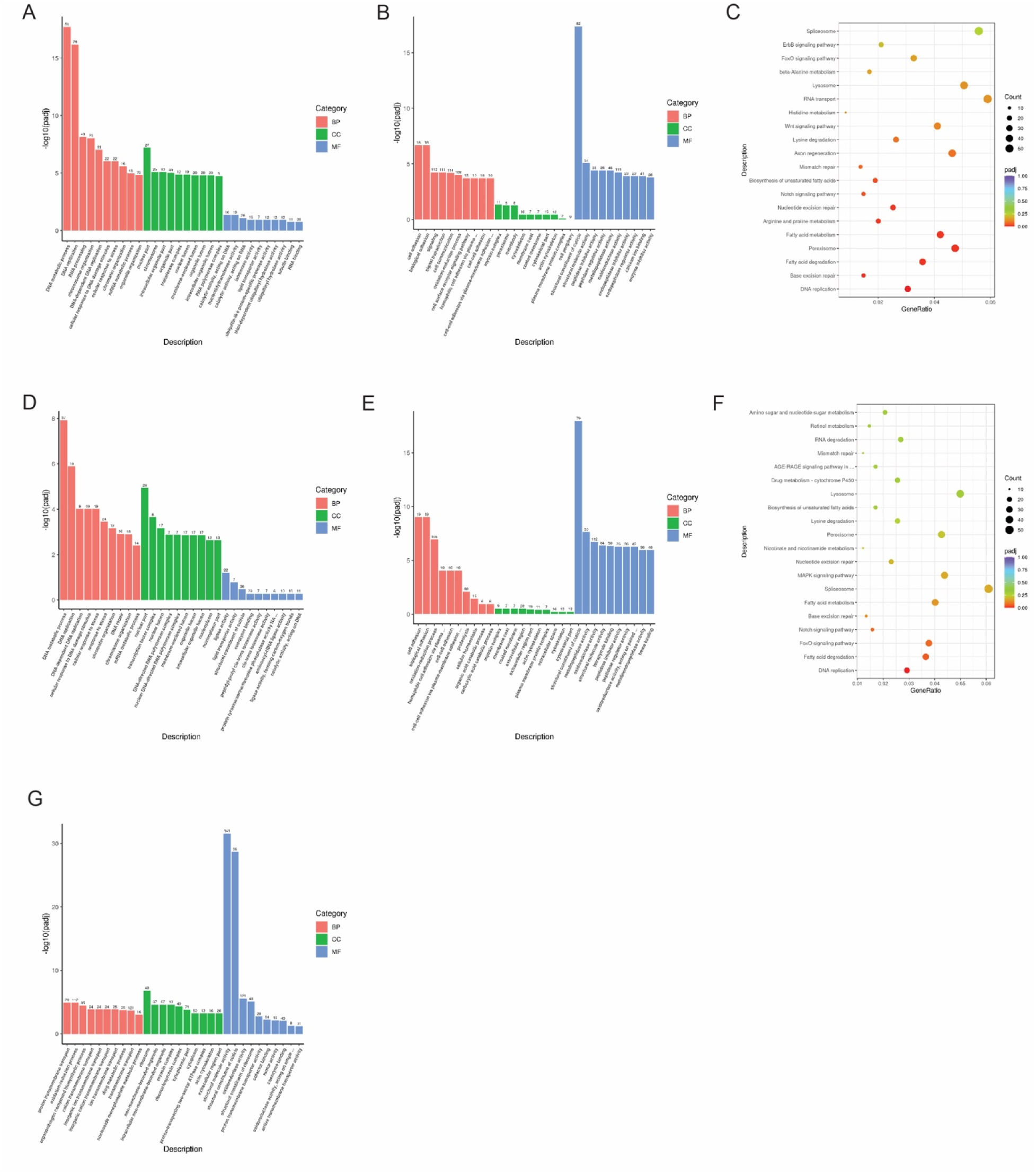
Different effects between OFA and OFB on healthspan promoting genes. Comparison of gene ontology (GO) and KEGG enrichment analysis between OFA vs. mock treated controls (**A-C**), OFB vs. mock treated controls (**D-F**), and OFA vs. OFB treatment. The mean expression level for each gene is indicated by log2FoldChange. All genes considered to be significant has adjusted p-value < 0.05.

**Figure S2.**
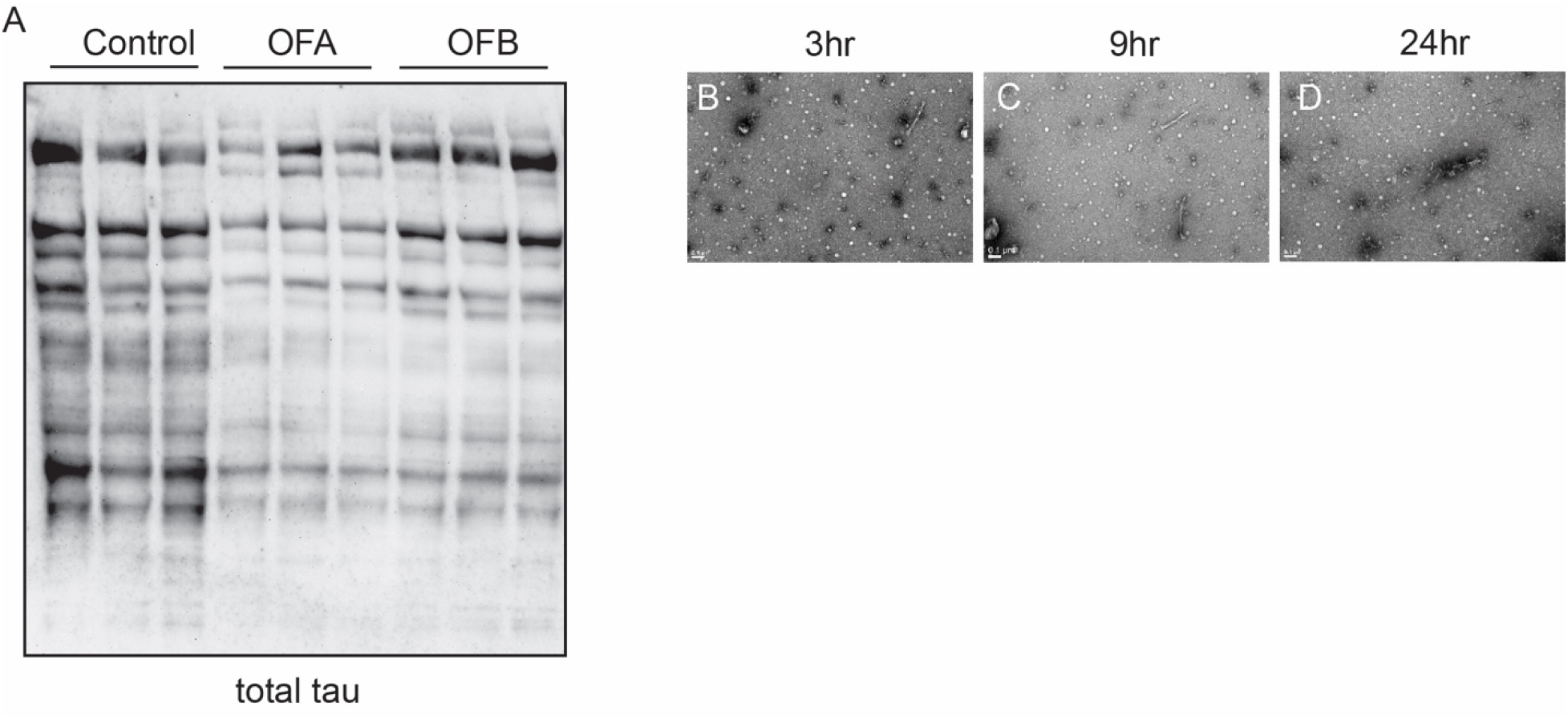
OF treatment reduces tauopathy across model systems. (**A**) Western blot analysis of total tau protein in hTau-expressing worms (KAE112) worms after treatment with OFA and OFB relative to mock-treatment (control). Additional representative images shown of AD tau fibrils, OFA condensates, and fibrils, after 3 hour (**B**), 9 hour (**C**), and 24 hours (**D**) of incubation with OFA.

**Figure S3.**
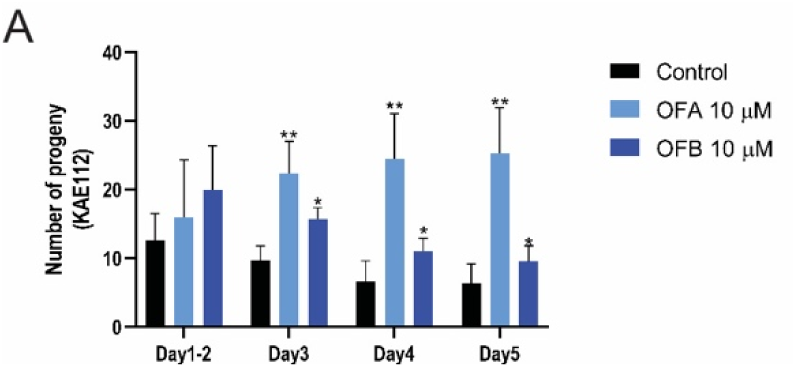
Effects of OFs on health metrics of in hTau-expressing animals. Comparison of daily progeny production between wild type (N2) and hTau-expressing worms (KAE112) worms treated with OFA and OFB as compared to mock treatment (control). *p<0.05, **p<0.01, ***p<0.001 and ****p<0.0001, compared to the mock-treated controls by one-way ANOVA following Bonferroni’s method (post hoc).

**Figure S4.**
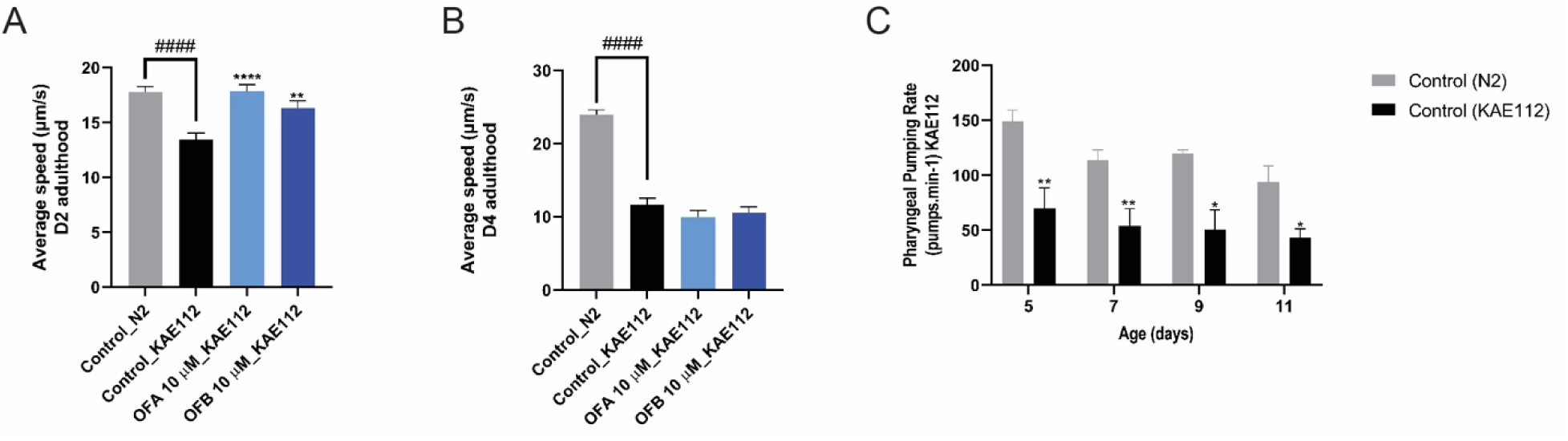
Effects of OFs on muscle functions of KAE112. Average crawling speed of wild type (N2) and hTau-expressing worms (KAE112) worms at D2 (**A**) and D4 (**B**) of adulthood after OFA and OFB treatment, as compared to mock treatment (control). Comparison of pharyngeal pumping rate between wild type (N2) and hTau-expressing worms (KAE112) worms (**C**). *p<0.05, **p<0.01, ***p<0.001 and ****p<0.0001, compared to the mock-treated controls by one-way ANOVA following Bonferroni’s method (post hoc).

**Figure S5.**
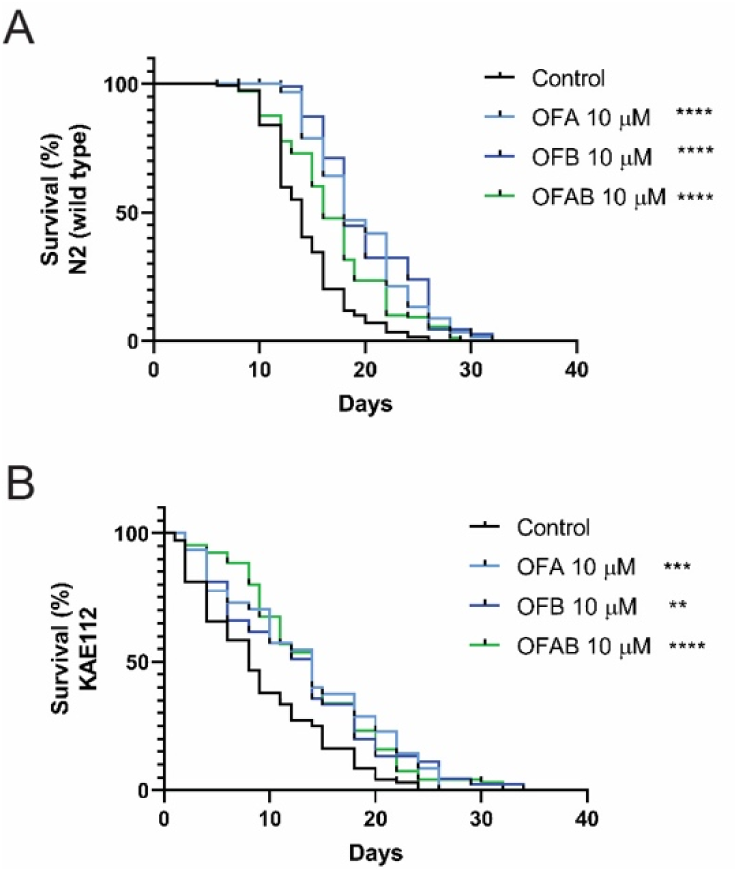
Effects of OFs on lifespan of KAE112. Survival curves of wild type (N2) (**A**) and hTau-expressing worms (KAE112) (**B**) worms treated with OFA, OFB, or OFA and OFB from L4. *p<0.05, **p<0.01, ***p<0.001 and ****p<0.0001, compared to the mock-treated controls by one-way ANOVA following log-rank test.

## Supplementary Table

Table S1. List of DEGs in OFA and OFB treatment compared to the untreated control in wild-type (N2) worms with the threshold of P≤0.05. The GO functional enrichment and KEGG pathway enrichment analyses of DEGs in OFA and OFB treatment compared to the untreated control in wild-type (N2) worms with the threshold of P≤0.05 are listed for each of the following GO-ontologies: Biological Process (BP), Cellular Component (CC), Molecular Function (MF).

## MATERIALS & METHODS

### *C. elegans* strains and maintenance

All strains were cultured on nematode growth media (NGM) supplemented with *Escherichia coli* OP50 using standard methods [36]. Worms were maintained at 20 °C. Strains used in this study include N2 Bristol wild type (N2), and KAE112 (*seals201 [myo-3p::human tau (0N4R;V337M)::unc-54 3’UTR* + *vha-6p::mCherry::unc-54 3’UTR*) [4]. Age-synchronized populations of worms were obtained by hypochlorite treatment [37].

### Oolonghomobisflavans

Oolonghomobisflavan A (OFA) (CAS No.126737-60-8, Cat No. NS240102) and oolonghomobisflavan B (OFB) (CAS No. 176107-91-8, Cat No. NS240202) were purchased from Nagara Science Co. (Gifu, Japan).

### Lifespan assay

Worms were synchronized to generate a synchronous L1 population. Larval stage 4 (L4) worms were moved to NGM agar plates supplemented with M9 buffer (untreated control) or OFs including 10 µM OFA, 10 µM OFB, or 5 µM OFA and 5 µM OFB (OFAB). The different concentrations of OFs were prepared in M9 buffer and placed above *E. coli* OP50 lawn and incubated at room temperature overnight before use. Animals were observed and moved to fresh medium every other day until the end of life. Worms that failed to respond to a gentle touch were scored as dead. Animals with internally hatched progeny, extruded gonads, or crawled to the side of the plate were censored. Each experimental replicate measured a minimum of 30 individual animals for a total of 90-120 animals/treatment.

### Pharyngeal pumping assay

Pharyngeal pumping assays and lifespan assays were conducted at the same time, specifically on the 5th, 7th, 9th and 11th h day of adulthood of wild-type (N2) and KAE112 worms. The pharyngeal pumping rates were quantified by counting pharynx contractions for 60s. Each experimental replicate measured a minimum of 20 individual animals for a total of 60-90 animals/treatment.

### Reproduction assays

Wild-type (N2) and KAE112 worms were synchronized in the same way as in the lifespan assay. The L4 larval stage animals were sorted and placed one by one on each NGM agar plate supplemented with OFs. For brood size assays, L4 worms were singled on NGM agar plate supplemented with oolong tea extracts and incubated at 20°C for 24 h. Each group had a minimum of 20 worms. The adult worms were moved every 12 h until egg-laying ceased. The eggs were counted using a dissecting microscope every day for 5 days to obtain a number of progeny and a mean brood size.

### WormLab measurement

Wild-type (N2) and KAE112 worms were synchronized by hypochlorite treatment. Eggs were allowed to hatch overnight for a synchronous L1 population on NGM agar plates supplemented with OFs. Worms were then allowed to grow until day 2 and day 4 of adulthood (day 2; 156 h from egg synchronization of wild-type (N2), 204 h from egg synchronization of KAE112, day 4; 204 h from egg synchronization of wild-type (N2), 252 h from egg synchronization of KAE112). At each time point, worms were washed with M9 buffer (+0.1% Triton) and dropped on an unseeded NGM plate. Worms were allowed to roam for 1 hour before recording crawling and thrashing with the MBF Bioscience WormLab microscope. The worms that moved at least 90% of the time were used to analyze with WormLab version 2022 software.

### Development assay

For body length measurements, wild-type (N2) and KAE112 worms were synchronized and treatment in the same way as described above. Worms were then allowed to grow until each time point (60 h, 108 h, and 156 h) and imaged by MBF Bioscience WormLab microscope. Body length measurements were quantified using WormLab version 2022 software.

### Paralysis assay

KAE112 worms were synchronized in the same way as in the lifespan assay and treated with OFs at the L4 stage on NGM agar plate. The number of paralyzed worms were counted from day 1 of adulthood. Worms were classified as paralyzed when they did not move or only moved their head (cleared bacteria giving a halo appearance around the worms’ heads). Paralyzed worms were recorded and excluded from the plates every other day.

### RNA sequencing

The L1 larval stage animals were treated with 10 µM OFA, 10 µM OFB, or M9 buffer (untreated control). After 48 h from treatment, L4 animals were washed with M9 buffer and frozen in TRI reagent at −80°C until use. Animals were homogenized and RNA extraction was conducted by using the Zymo Direct-zol RNA Miniprep kit (Cat No. R2052). Qubit^TM^ RNA BR Assay kit was used to determine RNA concentration. The RNA samples were sent to Novogene to perform RNA sequencing. Read counts were used for differential expression (DE) analysis by using the R package DEseq2 (R version 3.5.2). Differentiated expressed genes were analyzed using p value <0.05 and fold change >1.5 as cutoff.

### Western blot analysis

Synchronized populations of KAE112 worms were grown to the third day of adulthood. Worms were washed off plates with M9 buffer and fractured by freeze-thaw cycles in liquid nitrogen. The fractured worm biomass was grounded and lysed in FA buffer (1 mM EDTA pH 8.0, 0.1% w/v Sodium deoxycholate, 1% v/v Triton X-100, 1x HALT protease inhibitor). Total protein concentrations were quantified by Bradford assay (Sigma). An equal amount of protein (20 µg) was separate on 4%-12% bis-tris polyacrylamide gel (Invitrogen) in MOPS running buffer (Invitrogen,) and then transferred to nitrocellulose membranes (GE Healthcare Life science). After blocking for 1 h with 3% BSA in PBST (PBS, 0.1% Tween 20), the membranes were subjected to immunoblot analysis. Antibodies used include: P-tau S202 clone D4H7E (Cell Signaling, 1:1000), P-tau S416 clone D7U2P (Cell Signaling, 1:1000), pan-tau (Millipore Sigma, 1:1000), Histone H3 (Abcam, 1:5000), and HRP-conjugated secondary antibodies (Thermo Fisher, 1:10,000). Specific protein bands were visualized and evaluated using FluorChem HD2 (Protein Simple).

### Statistical analysis

Data are presented as mean ± SEM (n, indicated for each experiment, replicated a minimum of three times). Data were analyzed by one-way ANOVA following Bonferroni’s method (post hoc). Data handling and statistical processing were performed using GraphPad Prism 9.0. Differences were considered significant at the *p* ≤ 0.05 level.

### K18CY cell culture

HEK293T cell lines that stably express tau-K18CY labeled with green fluorescent protein (GFP) obtained from Marc Diamond’s laboratory at the University of Texas Southwestern Medical Center (Sanders *et al*., 2018) were used. The cells were cultured in a T25 flask in Dulbecco’s Modified Eagle Medium (DMEM) (Life Technologies, cat. 11965092) supplemented with 10% (vol/vol) Fetal Bovine Serum (FBS) (Life Technologies, cat. A3160401), 1% penicillin/streptomycin (Life Technologies, cat. 15140122), and 1% Glutamax (Life Technologies, cat. 35050061) at 37°C and 5% CO_2_ in a humidified incubator. To test the inhibitors on the biosensor cells, 100 µl of cells were plated in 96 well plates and stored in the 37°C, 5% CO_2_ incubator for 16 to 24 hours prior to transfection.

### Biosensor cell seeding assays

EGCG (control) and OFs were diluted in dimethyl sulfoxide (DMSO) to 1.4 mM stocks. Homogenized AD brain was diluted in Opti-MEM (Thermo Fisher Scientific, cat. 31985062) in a 1:20 ratio. Diluted brain homogenate was incubated with indicated inhibitors for 16 to 24 hours at 4°C. Inhibitor-treated seeds were sonicated again in a Cup Horn (Qsonica, MPH) water bath for 3 minutes at 40% power and then mixed with a 1 to 20 solution of Lipofectamine 2000 (Thermo Fisher Scientific, cat. 11668019) and Opti-MEM. The Lipofectamine creates a liposome around the fibrils to allow delivery into the cells. After 20 minutes, 10 µl of inhibitor-treated fibrils were added to the previously plated 100 µl of cells in triplicate, avoiding use of the perimeter wells to yield a final ligand concentration of 10 mM on cells.

### Preparation of crude Alzheimer’s brain-derived tau seeds

Human Alzheimer’s brain autopsy samples were obtained from the UCLA Pathology Department according to HHS regulation from patients who consented to autopsy. Approximately 0.2 g of tissue was excised, and a Kinematica PT 10-35 POLYTRON was used to homogenize the tissue with 0.75 ml sucrose buffer (0.8 M NaCl, 10% sucrose, 10 mM Tris– HCl, pH 7.4) supplemented with 1 mM ethylene glycol tetraacetic acid (EGTA) at level 4-5 in 15 ml falcon tubes. Homogenates were aliquoted and used for seeding as described. For qEM studies, tau PHFs were further purified from homogenates by sarkosyl extraction. Briefly 0.5-1.0 g homogenized tissue was centrifuged at 15,300 rpm for 20 min. The supernatant was adjusted to a final concentration of 1% sarkosyl and incubated for 1 hr at room temperature with shaking at 250 rpm. Fibrils were obtained by ultracentrifugation at 95,000 rpm for 1 hr. Pellets were resuspended in sucrose buffer supplemented with 1 mM EGTA and 5 mM Ethylenediaminetetraacetic acid (EDTA) and centrifuged once more at 15,300 rpm for 20 min followed (for the supernatant) by ultracentrifugation at 95,000 rpm for 1 hr. The final pellet was resuspended in 0.1 ml of 20 mM Tris–HCl pH 7.4, 100 mM NaCl.

### Negative stain grid preparation

Purified Alzheimer’s brain-derived tau PHF fibrils were diluted 1:10 in PBS and incubated with OFA for indicated timepoints at 4°C. For qEM, after 48 hours incubation, EM grids were prepared by depositing 6μl of samples on formvar/carbon-coated copper grids (400 mesh) for 3 minutes with inhibitor pre-incubation times of either 0 hours (negative control) or 48 hours (positive control). The sample was rapidly and carefully removed by fast blot using filter paper without drying the grid and stained with 4% uranyl acetate for 2 minutes, then wicked dry by filter paper. Automated images were collected using the FEI Glacios driven by EPU software. Visible fibrils were counted from 66 images, each for the 0 and 24 hr OFA incubation time points, and fibrils were plotted by dividing counted images in thirds to evaluate standard error.

## ACKNOWLEDGMENTS

We thank S. Ledgerwood for technical assistance and C.M. Ramos for critical reading of the manuscript. This work was funded by the NIH R01AG058610 and Hevolution Foundation award HF AGE-004 to SPC, and an AFAR Postdoctoral fellowship to CD, and a pilot grant to PMS that is funded in part by the Nathan Shock Center of Excellence P30AG068345. We also thank the USC School of Gerontology Imaging Core. Some strains were provided by the CGC, which is funded by the NIH Office of Research Infrastructure Programs (P40 OD010440). We thank WormBase for database curation and data access.

## Author contributions

Conceptualization: PMS and SPC; Methodology: CD, XC, PMS, and SPC; Investigation: CD, XC, PMS, and SPC; Visualization: CD, XC, PMS, and SPC; Supervision: SPC; Writing (original draft): CD and SPC; Writing (reviewing & editing): CD, XC, PMS, and SPC.

## Data and materials availability

All data are available in the main text or the supplementary materials. Sequencing data is available at GEO.

